# Complete genomic characterisation of two *Escherichia coli* lineages responsible for a cluster of carbapenem resistant infections in a Chinese hospital

**DOI:** 10.1101/100941

**Authors:** Zhiyong Zong, Samuel Fenn, Christopher Connor, Yu Feng, Alan McNally

**Author notes:** Corresponding author: Dr Alan McNally, Institute of Microbiology and Infection, College of Medical and Dental Science, University of Birmingham, Birmingham B15 2TT. 0044 121 4158433.

## Abstract

The increase in infections as a result of multi-drug resistant strains of *Escherichia coli* is a global health crisis. The emergence of globally disseminated lineages of *E. coli* carrying ESBL genes has been well characterised. An increase in strains producing carbapenemase enzymes and mobile colistin resistance is now being reported, but to date there is little genomic characterisation of such strains. Routine screening of patients within an ICU of West China Hospital identified a number of *E. coli* carrying the *bla*_NDM-5_ carbapenemase gene, found to be two distinct clones, *E. coli* ST167 and ST617. Interrogation of publically available data shows isolation of ESBL and carbapenem resistant strains of both lineages from clinical cases across the world. Further analysis of a large collection of publically available genomes shows that ST167 and ST617 have emerged in distinct patterns from the ST10 clonal complex of *E. coli*, but share evolutionary events involving switches in LPS genetics, intergenic regions and anaerobic metabolism loci. These may be evolutionary events which underpin the emergence of carbapenem resistance plasmid carriage in *E. coli*.

## Background

Infections from multi-drug resistant (MDR) *Escherichia coli* are a significant global health care threat[1]. Despite being an extremely diverse species, MDR in *E. coli* is largely confined to strains capable of causing extra-intestinal infections (ExPEC) such as urinary tract infections (UTI) and bacteraemia[1-4]. As many as 50% of *E. coli* strains isolated from UTI and bacteraemia cases may exhibit resistance to three or more classes of antibiotic, termed MDR. This resistance is primarily driven by the acquisition of large plasmids containing multiple resistance genes[2]. The rapid global dissemination of MDR *E. coli* is associated with carriage of plasmids containing genes encoding extended-spectrum β-lactamases (ESBL) which confer resistance to third-generation cephalosporins[5]. The carriage of MDR plasmids containing ESBL genes renders *E. coli* susceptible only to the carbapenem class of antibiotics and the antimicrobial compound colistin[5]. However strains of *E. coli* are now being reported with plasmids containing β-lactamases conferring resistance to carbapenems (carbapenemases) and the *mcr-1* colistin resistance gene [6-9]. The global dissemination of ESBL *E. coli* is attributable to the rapid dispersal of a small number of *E. coli* lineages. The most dominant of these is the ST131 lineage which is predominantly associated with carriage of the *bla*_CTX-M-15_ ESBL gene[2]. ST131 is an ExPEC lineage and the most common cause of UTI and bacteraemia in the developed world[2]. Other dominant lineages of ESBL *E. coli* are ST73, ST95, and ST648 which are also ExPEC[3,4]. ESBL carriage can also be found transiently in strains belonging the ST10 clonal complex of *E. coli*[3]. ST10 complex strains are host generalist *E. coli* which are frequently found as intestinal commensal inhabitants of mammals and avian species[10], and are devoid of the virulence-associated genes known to be required for pathogenesis[11]. Our knowledge of the genomic landscape of carbenemase production in *E. coli* is far less developed, with the vast majority of reports being genomes of individual clinical isolates sporadically distributed across the globe. Just one significant publication exists reporting a specifically designed genomic analysis of a temporal collection of carbapenem resistant *E. coli* which showed very wide dissemination of carbepenem resistance across species and within-species lineages of the enterobacteriaceae [12]. Here we report the isolation of *E. coli* containing the carbapenem-resistance gene *bla*_NDM-5_ in an ICU ward in West China Hospital, Chengdu. The isolates do not belong to one of the dominant MDR lineages of ExPEC, but to ST167 and ST617, both members of the ST10 clonal complex. Genomic data supports the long-term presence of these bacteria in the ICU with repeated dissemination from a central reservoir. Contextualisation of the Chinese strains with a collection of publically available genomes shows isolation of MDR ST167 and ST617 strains from clinical episodes across the world, and in the case of ST167 frequent occurrence of carriage of both ESBL and carbapenemase genes. By comparing these lineages to a large number of publically available ST10 genomes we identify potentially significant events in their evolutionary trajectories, including mutations in the LPS biosynthesis locus which truncate LPS. We also find evidence of compensatory mutations in intergenic regions as found in *E. coli* ST131 as well as mutations in anaerobic metabolism loci. Our findings support the need for a more concerted global surveillance effort focussing on identifying frequently occurring lineages of carbapenem resistant *E. coli*.

## Methods

### Bacterial isolation and characterisation

Strain 0215 was recovered from a rectal swab of a 75-year-old male patient on September 2013 in a 50-bed medical ICU at West China Hospital, Chengdu, during routine screening that is performed as standard in the ICU on all new admissions. Following the identification of *bla*_NDM-5_, we performed an active screening project on adult patients (age ≥16) at the medical ICU ward during a 7-month period from May to November 2014. This study was conducted in accordance with the amended Declaration of Helsinki and was approved, under a waiver of consent, by the Ethics Committee of West China Hospital. Rectal swabs were collected from patients within 2 days of admission to the ICU and within the 3 days prior to ICU discharge for those patients with a length of stay of 3 days or more. Swabs were transferred to the laboratory in transport media and were screened for carbapenem-resistant Enterobacteriaceae using the CHROMAgar Orientation agar plates containing 2 μg/ml meropenem. Carbapenem-resistant *E. coli* were recovered from the rectal swabs of 8 different patients (Table 1). Furthermore, one of the 8 patients developed bacteraemia during his ICU stay and an *E. coli* was recovered from his blood and included in the study. During the study period, two additional *E. coli* clinical isolates carrying *bla*_NDM-5_ were recovered in the hospital, from two patients on admission.

**Table 1.** Sources and patients of E. coli isolated in West China Hospital carrying bla_NDM-5_

| Isolate | ST | Collection date | Collection, days after admission to ICU | Source | The host patient |  |
| --- | --- | --- | --- | --- | --- | --- |
|  |  |  |  |  | Age | Sex |
| 0215 | 167 | 2013-09 | 0 | Rectal swab | 75 | Male |
| 243 | 167 | 2014-05 | 0 | Rectal swab | 84 | Female |
| 442<sup>a</sup> | 167 | 2014-07 | 7 | Rectal swab | 39 | Male |
| 57<sup>a</sup> | 167 | 2014-07 | 16 | Blood | 39 | Male |
| 936 | 167 | 2014-09 | 12 | Rectal swab | 63 | Female |
| 1222 | 167 | 2014-10 | 7 | Rectal swab | 17 | Male |
| 1237 | 167 | 2014-10 | 3 | Rectal swab | 44 | Female |
| 25 | 167 | 2014-10 | 0 | Ascite | 45 | Female |
| 784 | 617 | 2014-08 | 0 | Rectal swab | 82 | Male |
| 1037 | 617 | 2014-09 | 12 | Rectal swab | 85 | Male |

<sup>a</sup>Isolates 442 and 57 were recovered from the same patient.

### Genome sequencing

The ST167 and ST617 strains isolated in Chengdu were cultured in LB broth at 37°C overnight. DNA was extracted using QIAamp^®^ DNA Mini Kit (QIAGEN) and 150 bp paired-end libraries of each strain prepared and sequenced using the Illumina HiSeq X-Ten platform (raw data accession numbers Table S1 and S2). Genomes were assembled using SPAdes[13] and annotated using Prokka[14]. The MLST sequence type of the strains was determined using the in silico prediction tool MLSTFinder[15]. The *E. coli* genome database Enterobase (www.enterobase.warwick.ac.uk) was interrogated on 1^st^ December 2016 and all available ST167 and ST617 genomes were downloaded (Table S1 and S2) and annotated using Prokka. A further 256 ST10 genomes were selected to represent the geographical, temporal, and source attribution diversity present in the database (Table S3) and were downloaded and annotated using Prokka. To select these genomes a phylogenetic tree was inferred from the assembled genome of every ST10 on Enterobase using Parsnp[16]. From this phylogeny 500 genomes were chosen to span the entire phylogenetic diversity, and then the final selection made to represent the full ST10 diversity as described. The antibiotic resistance gene profile of all isolates was determined using Abricate (https://github.com/tseemann/abricate).

### High-resolution SNP analysis

We created a closed genome sequence for a Chinese ST167 strain 1237 by combining our Illumina sequence data with data generated on the MinIon sequencer. Raw MinIon reads were converted into fastQ format (accession number PRJNA422975) using Poretools [17] and assembled using Canu [18], resulting in a single contig chromosome and four distinct single contig plasmids. The raw illumina data was then used to polish the genome assembly via five iterative rounds of polishing with Pilon [19]. The ST167 and ST617 genomes from Chengdu were analysed by mapping raw reads against the hybrid assembled ST167 genome. Mapping was performed using Snippy (https://github.com/tseemann/snippy) and the resulting SNP profiles were used to create a consensus sequence for each genome which was aligned using the parsnp alignment tool in Harvest[16]. Analysis of the plasmid containing the *bla*_*NDM-5*_ gene revealed that it was a 47-kb IncX3 plasmid and there were no antibiotic resistant genes other than *bla*_NDM-5_ located on the plasmid. Specific mapping of the raw Illumina data against the pNDM5 plasmid was performed for all strains as described above.

### Phylogenetic analysis

Pan-genomes were constructed for the ST167, ST617, ST10, and combined datasets using Roary[20] with the --e --mafft setting to create a concatenated alignment of core CDS. The alignments were used to infer ST167, ST617, ST10, and combined phylogenies using RaxML[21] with the GTR-Gamma model of site heterogeneity and 100 bootstrap iterations. Carriage of ESBL and carbapenemase genes was annotated on the trees using Phandango (https://jameshadfield.github.io/phandango/), and geographical source was annotated using iTOL[22].

### Detection of lineage specific genetic traits

Microbial GWAS was performed using two approaches. First the combined data set pan-genome matrix was used as input for Scoary [23] searching for loci unique to ST167, ST617, and both ST167 and ST617 versus ST10. In parallel we also used SEER [24] to detect kmers significantly associated with ST167, ST617, or both combined versus ST10. The results of both approaches were combined to identify coding loci associated with the emergence of ST167 and ST617. In silico serotyping was performed using two independent methods, SRST2 and SerotypeFinder [25,26]. Both methods utilise WGS data to specific O and H antigens to strains. Intergenic regions (IGRs) were investigated using Piggy [27] to search for IGRs which had switched [28] in ST617, ST167, or both compared to ST10. This data was combined with SEER data to identify high-confidence IGR switches associated with the emergence of ST167 and ST617.

## Results

### Presence of *E. coli* ST167 and ST617 strains containing the NDM-5 carbapenemase resistance gene in an ICU ward in West China Hospital

A total of ten isolates of *E. coli* containing *bla*_NDM-5_ were obtained during the investigation. Nine of these isolates belonged to sequence types ST167/617 (Table 1), which are members of the ST10 complex of *E. coli* most commonly associated with mammalian intestinal commensal carriage. Three ST167 isolates (0215, 243 and 25) were obtained from swabs or clinical samples collected on admission to hospital, suggesting that they were introduced from external sources. The three patients were all citizens of Chengdu city but they were admitted to different local hospitals before transferring to West China hospital. The remaining ST167 isolates were recovered from swabs or samples collected at least 3 days after admission to the ICU of West China hospital, suggesting that they were acquired during their ICU stay. ST167 *E. coli* carrying *bla*_NDM-5_ caused infections (bacteremia and abdominal infection) in only two patients but colonised the others. Both ST617 *E. coli* carrying *bla*_NDM-5_ only colonised patients. All patients colonised or infected with *E. coli* carrying *bla*_NDM-5_ of ST167 or ST617 had received carbapenems before the recovery of the isolates.

### SNP analysis suggests continued dissemination of strains from a central reservoir and sharing of resistance plasmid between lineages

To determine the level of relatedness between all isolated strains we mapped reads of all the strains against a closed ST167 strain (strain 1237) generated by a combination of Illumina and MinIon sequence data. The resulting high-resolution SNP alignment showed the distance between the ST167 and ST617 strains to be over 25,000 SNPs, confirming they are distinct lineages, with the two ST617 isolates separated by just 7 SNPs. Deeper analysis of the ST167 cluster of strains showed diversity ranging from 5 to 799 SNPs (Fig 1). Strains 936 and 1222 (both carriage isolates) are the most closely related isolates with just 5 SNPs difference between them, with both strains being acquired by patients in the ICU within one month of each other. However these strains are 73 SNPs different from a strain isolated the exact same month on the ICU from a strain (1237) that was acquired in the ICU. This is almost double the genetic distance (46 SNPs) from a strain acquired (442 and 57, isolated from the same patient) in the ICU two months earlier. These distances are also larger than those for any isolate to the first two strains brought into the ICU, strain 0215 and strain 243, which differ from all other isolates by around 30 SNPs, and from each other by 15 SNPs. Such an observation suggests a potential combination of patient-to-patient transmission in the affected ICU [29], along with the continued dissemination of the strain from a central reservoir where there is an accumulation of diversity [29,30]. Genomic analysis also allows us to identify a second introgression of an ST167 strain (25) from the community, which is over 700 SNPs different from the other isolates. Mapping of the raw sequence data against the 43kb IncX3 plasmid containing *bla*_*NDM-5*_ also confirmed that the plasmid present in the ST617 strains was identical to that in all of the ST167 strains with just two detectable SNPs difference across the isolates.

**Figure 1:**
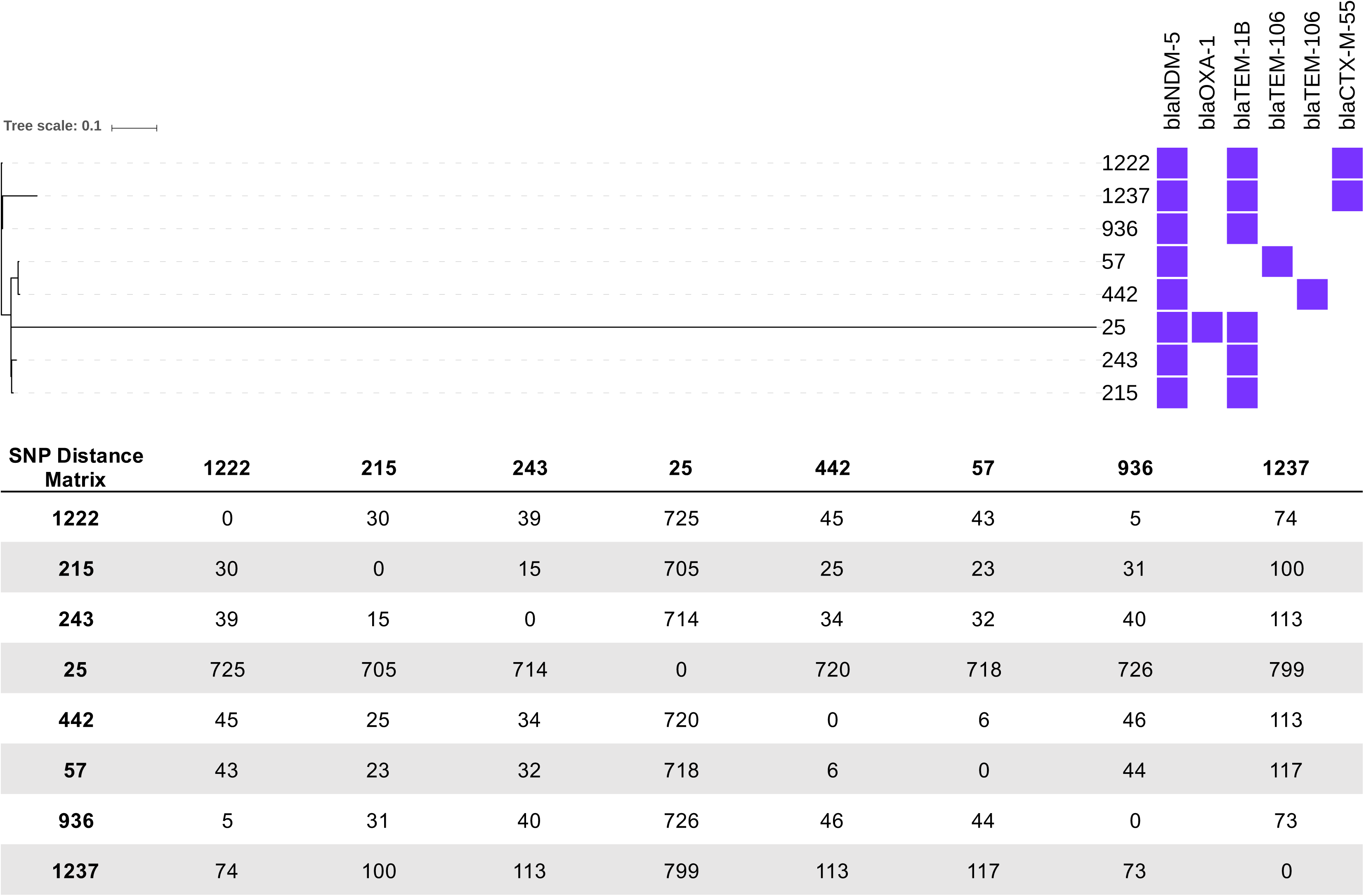
Maximum likelihood phylogenetic tree of *E. coli* ST167 strains isolated form the ICU of West China hospital. The phylogeny is inferred from a SNP aligment obtained by mapping raw data against a MinIon/Illumina hybrid complete assembly of isolate 1237. The annotation denotes the presence of ESBL and CPE associated β-lactamases as determined by Abricate.

### MDR ST167 and ST617 *E. coli* have been isolated across the world

We sought to contextualise the wider relevance of our Chengdu isolates by investigating the wider prevalence of ST167 and ST617 strains. We searched the Enterobase *E. coli* database and recovered a total of 87 genomes of ST167 (table S1) and 86 genomes of ST617 (table S2), isolated from across the world. A core CDS-based phylogeny of both lineages showed a diverse set of genomes with around 17,000 SNPs in ST167 and around 15,000 SNPs in ST617. Annotation of the ST617 phylogeny with β-lactamase gene carriage shows a high prevalence of the *bla*_CTX-M-15_ ESBL gene in characterised isolates (Fig 2A). Annotation of the ST167 phylogeny with β-lactamase gene carriage (Fig 2B) shows a pattern of resistance gene carriage, with multiple independent acquisitions of carbapenemase across the phylogeny including *bla*_NDM-1_, *bla*_NDM-5_, *bla*_NDM-7_, *bla*_OXA-181_, and *bla*_KPC-3_. For both phylogenies there is clear evidence of isolation of strains from across the globe.

**Figure 2:**
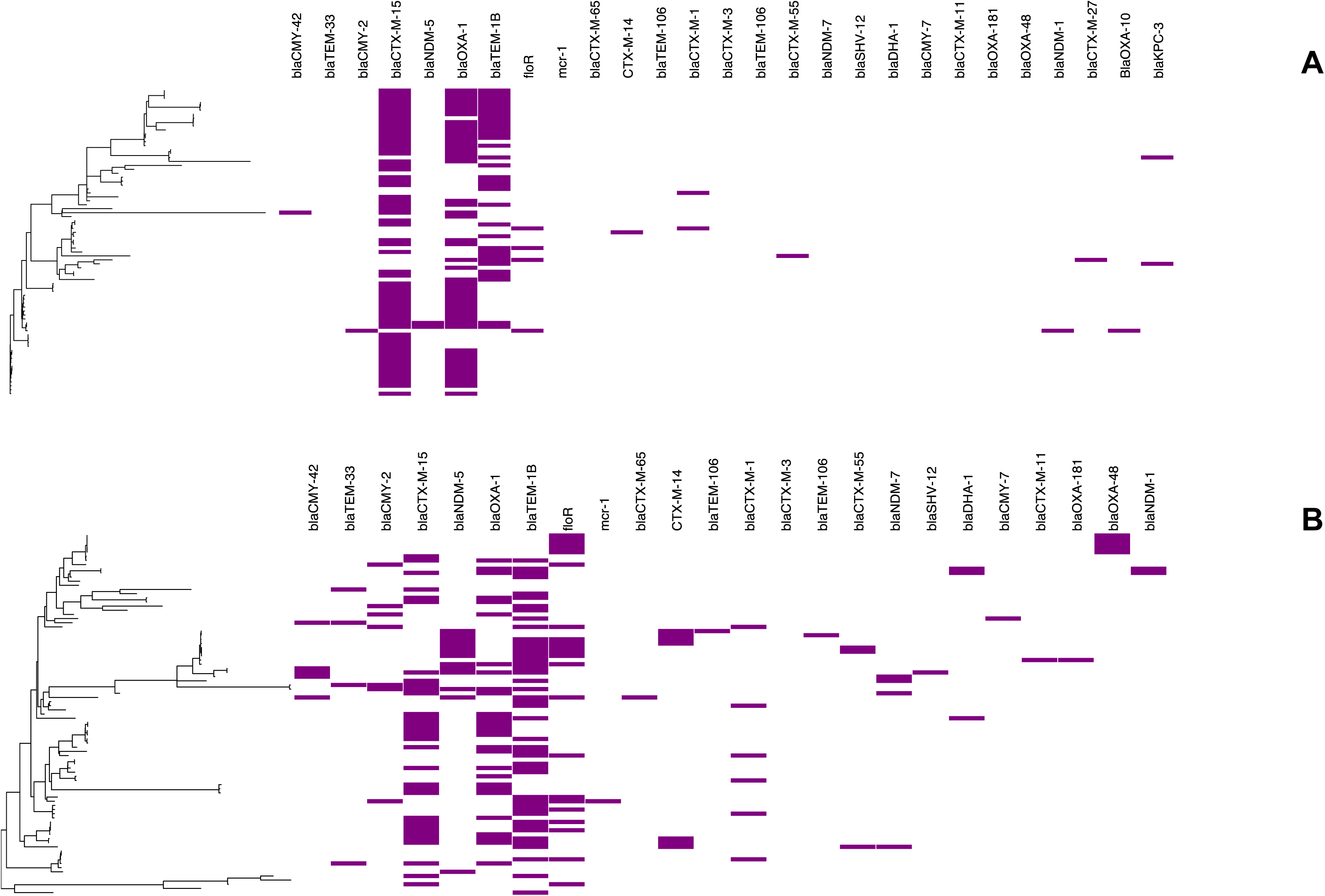
Maximum likelihood phylogenetic trees of a global collection of (A) ST617 and (B) ST167 strains. The phylogeny is inferred from an alignment of concatenated core CDS sequences as determined by Roary, and is mid-point rooted. The annotation denotes the presence of ESBL and CPE associated β-lactamases as determined by Abricate.

### Evolutionary genomic analysis correlates switches in LPS gene content with the emergence of the ST167/ST617 lineage

Both ST167 and ST617 are single locus variants of the ST10 lineage of *E. coli*. ST10 is the most abundant lineage of *E. coli* represented in the Enterobase database and contains isolates ranging from drug susceptible environmental and human commensal strains, to multi-drug resistant strains isolated from human clinical UTI and bacteraemia infections. We selected 256 ST10 genomes from Enterobase (Table S3) to represent the known spectrum of ST10 diversity present in the database, and merged this data set with our publically available ST167/ST617 genome data set to create a larger ST10 complex phylogeny (Fig 3). The resulting phylogeny shows that ST167 and ST617 are sister clades with respect to ST10, with ST617 emerging as a nested clade from a single outlying ST167 genome, though the distance between ST167 and ST617 is around 18,000 SNPs. Given the phylogenetic pattern of ST167 and ST617 with respect to ST10, we sought to determine if their emergence from ST10 is associated with defined evolutionary events. We used a combined GWAS approach to compare the ST167/617 genomes with ST10, using both SEER and SCOARY analysis of a pangenome matrix. Only loci considered to be significantly associated with one lineage over the other by both methods were further investigated (Dataset S1). Most striking was the absence of the *wzzB* gene and *wca* biosynthetic cluster in ST167/ST617 whilst the majority of the ST10 genomes contained both (Figure 4). These genes are involved in LPS biosynthesis with *wzzB* being the master controller of O antigen chain length in the *wzx/wzy* pathway, whilst *wca* genes are responsible for colonic acid biosynthesis [31]. In silico *E. coli* serotyping [32] established that ST167 and ST617 demonstrate the exact same O antigenic type (O32novel) with similarity also seen in H antigen type (H9 or H10) (Figure 3), whilst the SerotypeFinder database identified the strains as O89.

**Figure 3:**
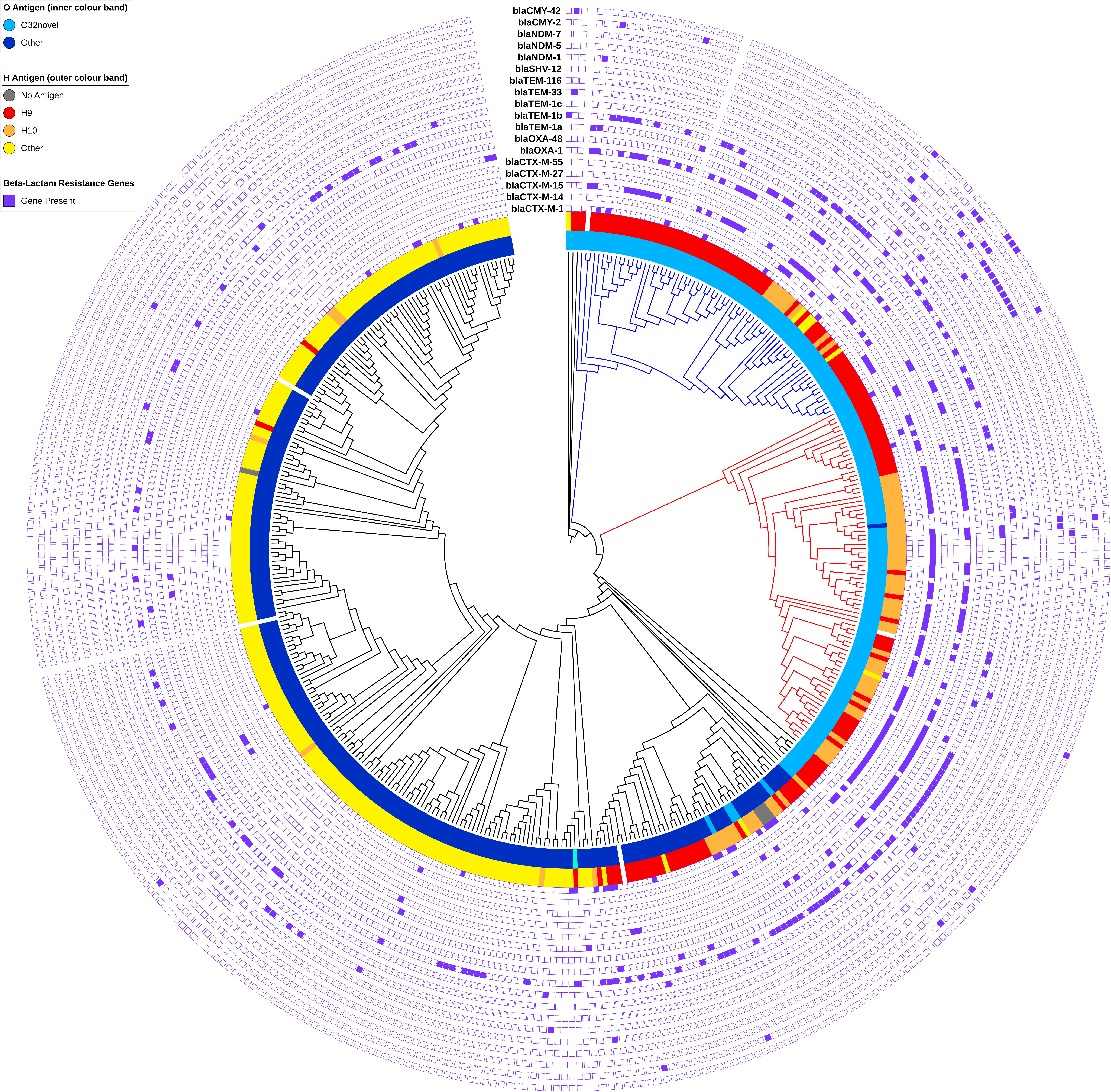
Cladogram inferred from a maximum likelihood phylogenetic tree of ST10 strains (black branches), ST617 (red branches) and ST167 (blue branches) strains. The phylogeny is inferred from an alignment of concatenated core CDS sequences as determined by Roary, and is mid-point rooted. The outermost annotation denotes the presence of ESBL and CPE associated β-lactamases as determined by Abricate. The inner ring of annotation on the tree indicates O-antigen type as determined by in silico typing using srst2. The outer ring of annotation indicates H-antigen flagellar type as determined in silico using srst2.

**Figure 4:**
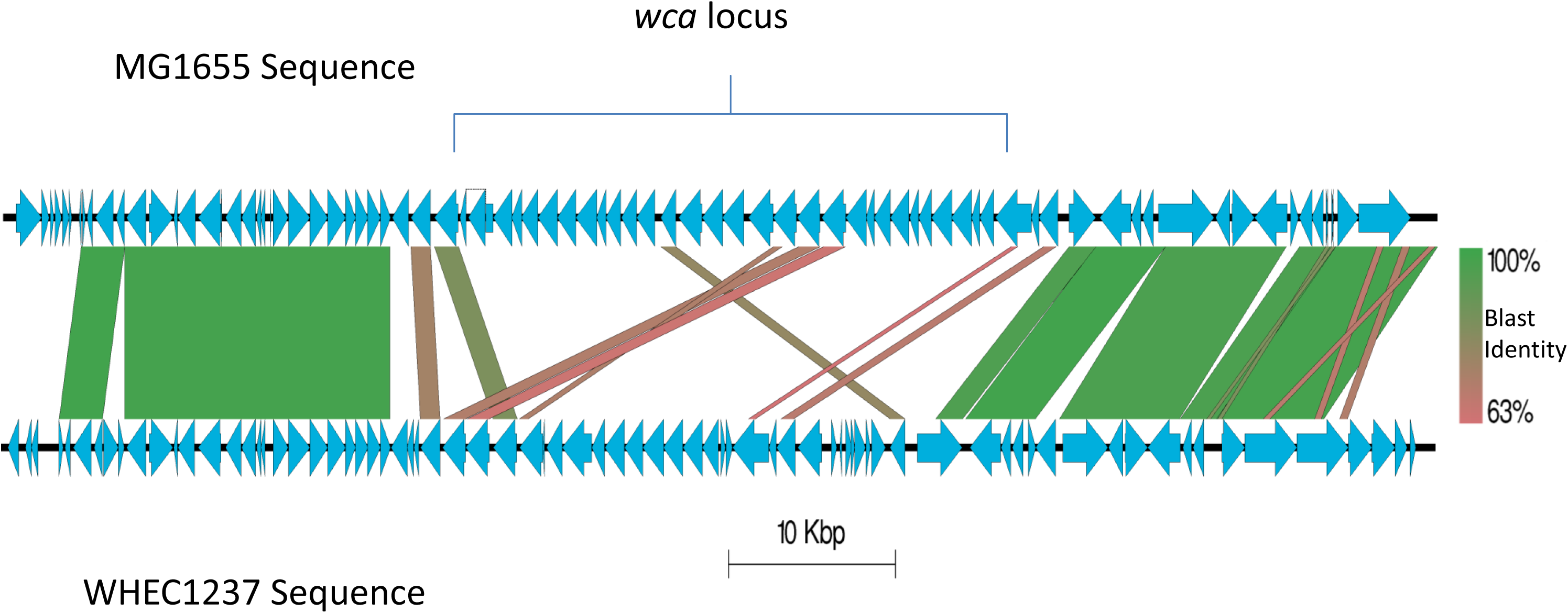
Diagrammatic comparison of the region of the genome of *E. coli* MG1655 (top of diagram) and the ST167 strain 1237 (bottom of diagram) containing the *wca* colanic acid biosynthesis locus.

Our combined GWAS analysis also identified another ~90 CDS which were present across the entire data set, but which had distinct alleles in the ST167/ST617 genomes compared to those in ST10 (Figure 5, Dataset S2). Many of these CDS encode dehydrogenase enzymes involved in anaerobic metabolism, or are part of the *cob/pdu/eut* operons known to be involved in anaerobic respiration during intestinal inflammation [33]. This would appear to suggest differential evolutionary events in key genes involved in anaerobic metabolism in the formation of the ST167/ST617 lineage. Also present were unique alleles in core CDS involved in acid and bile salt tolerance, and a number of fimbrial-like proteins. In conjunction these data would suggest differential evolutionary forces acting on loci involved in mammalian colonisation in ST167/617 in comparison to ST10. Furthermore a combined SEER and Piggy approach identified unique sequences in 17 intergenic regions (IGRs) upstream of core CDS in ST167/617 that were distinct from ST10, including IGRs upstream of anaerobic metabolic loci also present in the SEER/SCOARY analysis (Dataset S1).

**Figure 5:**
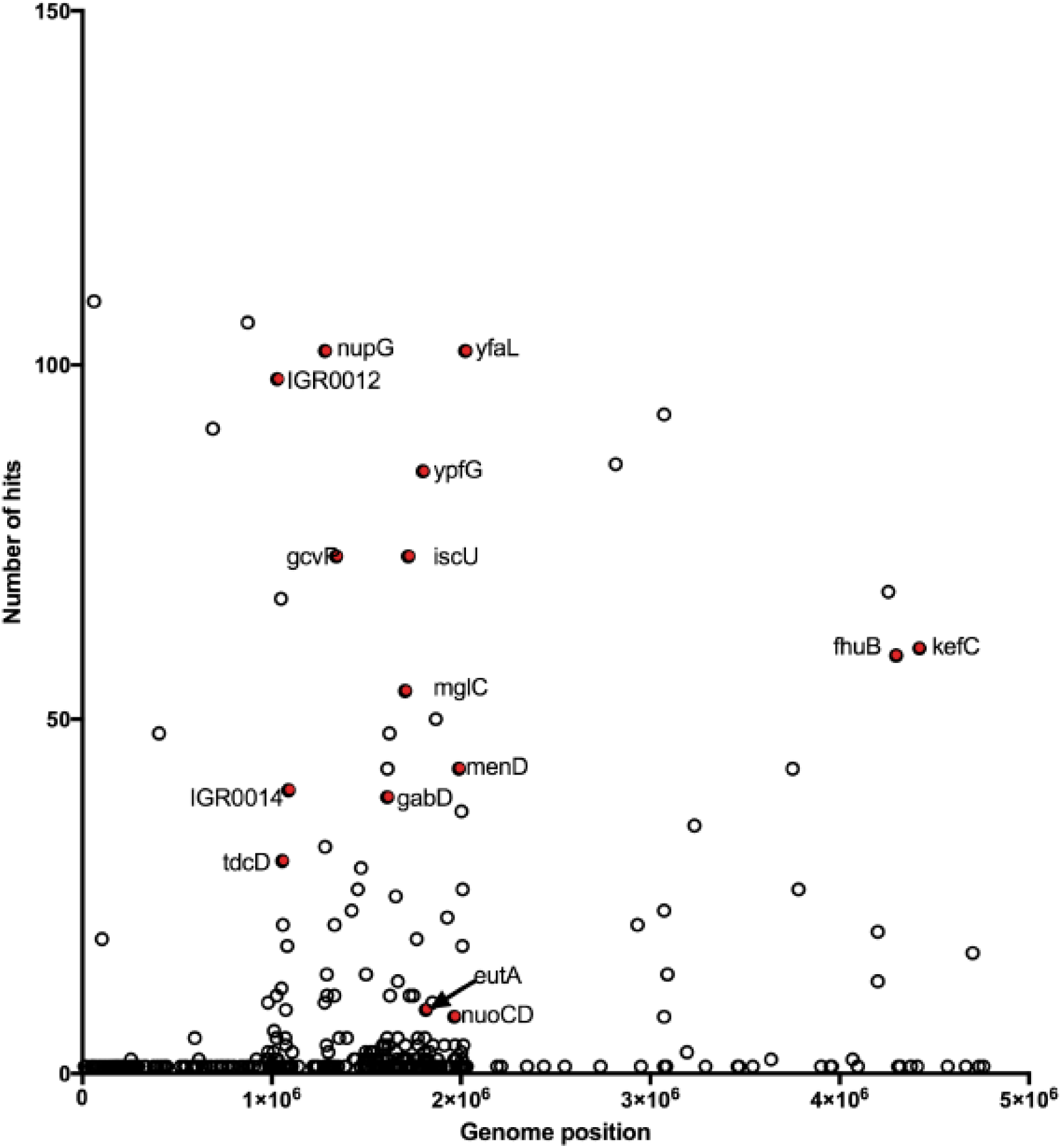
Manhattan skyline plot showing position of kmers identified by GWAS analysis as being significantly associated with ST167/617 compared to ST10. The x axis indicates the position on the WCHEC1237 complete genome assembly, whilst the Y axis indicates the numbers of statistically significant kmers mapping at that position. Hits indicated in red are either intergenic regions (labelled IGR) identified as being unique by both Piggy and SEER analysis, or anaerobic metabolism loci identified as significantly different by both SEER and Scoary.

## Discussion

Our data presented here provide a comprehensive genomic analysis of two lineages of carbapenem resistant *E. coli* infecting multiple patients within the ICU of West China hospital. Both these lineages, ST167 and ST617, are members of the larger ST10 complex of *E. coli*, which is ubiquitously found in environmental, human clinical, and mammalian intestinal commensal sampling. Our analysis is the first genome level characterisation of strains belonging to ST167 or ST617, despite a number of single site reports of clinical infections with both lineages existing in the literature. From a public health perspective our genomic characterisation of the ST167 and ST617 strains isolated form the ICU provide insight into the dynamics of carbapenem resistant *E. coli* infection. Our genomic epidemiology analysis of the ST167 strains suggests a scenario whereby a strain circulating in the Chengdu area enters the hospital setting and establishes a reservoir in the hospital environment, leading to continued episodes of acquisition and infection from a central source where diversity is accumulating [30]. This is also supported by the observation of ST617 being introduced into the ICU by a patient followed by acquisition in the ICU a month later by a strain just 7 SNPs different.

Our analysis also shows that the diversity which accumulates in the genome of the ST167 isolates during the course of the investigation is not mirrored by diversity in the plasmid carrying the *bla*_*NDM-5*_ gene. Only 1 SNP difference existed between the sequence of this plasmid in the ST167 isolates, and only 2 SNPs difference between the ST167 and ST617 isolates. As a result it is impossible to tell if the IncX3 plasmid associated with dissemination of *bla*_*NDM-5*_ in China [34] was transferred between ST167 and ST617 in the hospital, or if the plasmid is highly stable with only deleterious mutations occurring and quickly purged from the population. Clearly there is a need for more thorough and detailed analysis of various resistance plasmids within and between hospitals, such as was done recently for NDM-1 plasmids in Latin America [35].

The lack of appropriately designed isolate collection and sequencing strategy means it is impossible to conduct any form of genomic epidemiological analyses of these *E. coli* lineages beyond our Chinese investigation. However the ready availability of a large number of good-quality, curated genome assemblies in the Enterobase genome database do allow us to delve deeper into the evolutionary history of *E. coli* ST167 and ST617. Whilst data generated and uploaded to Enterobase is prone to a bias towards clinical MDR strains, it is still clear that ESBL and carbapenem resistant strains of both these lineages have been isolated from across the world over the past 20 or so years (Tables S1 and S2). Phylogenetic analysis of almost 200 publically available genome sequences, contextualised by an equal number of ST10 genomes allows us to determine that ST617 shares a common ancestor with ST167 distinct from ST10.

Comparative genomic analysis and GWAS for traits specific to ST167 and ST617 compared to ST10 also support emergence along a shared evolutionary branch. Key among these is the complete loss of the *wca* operon encoding colanic acid biosynthesis in the LPS biosynthesis pathway. The majority of *E. coli* produce their LPS utilising the O-unit translocation pathway encoded for by *wzx* and *wzy*[31]. This method utilises glycosyltransferases to assemble the O antigen in units at the cytoplasmic membrane. These units are then translocated by Wzx and polymerized by Wzy until the O antigen chain length is reached. This mechanism is utilised by the majority of the ST10 isolates, however genomic analysis shows that ST167 and ST617 utilise an alternative *wzm/wzt* ATP transporter pathway. This biosynthetic pathway assembles the entire O-antigen on the cytoplasmic face before Wzt transports the O-chain across [31], resulting in an O-antigen with truncated chain length. O-antigen chain length plays a major role in pathogenicity of Gram negative organisms, and it has been demonstrated that loss of long O-antigen chains in *Salmonella* optimizes immune evasion and allows successful colonisation [36]. Alongside the LPS genetic changes, we also observed unique alleles of anaerobic metabolism genes and genes potentially involved in host colonisation in ST167/617 compared to ST10. Recent modelling data has shown that any factor influencing the ability of a bacterium to colonise a host will also influence its likelihood of evolving antimicrobial resistance [37].

## Conclusions

We provide data for the first ever, single hospital genomic analysis of clinical isolates of carbapenem resistant *E. coli* belonging to the ST167/617 lineage. Our data presented here provide evidence for evolutionary events that would affect microbial interaction with a mammalian host underpinning the emergence of the ST167/617 lineage from ST10. There is also evidence for lineage specific alterations in intergenic regions in ST167/617, a phenomenon which has already been described as underpinning the emergence of MDR plasmid-containing *E. coli* ST131 strains [28]. Clearly there is now a need for a fully designed genomic epidemiological investigation of lineages of *E. coli* associated with carriage of carbapenem resistance plasmids arising from the ST10 clade. Such a study will fully inform us of any potential parallelism in the evolution of MDR lineages of *E. coli*, and of the true nature and scope of their prevalence and global dissemination.

## Declarations

### Ethics approval

Not applicable

### Consent for publication

Not applicable

### Availability of data

All raw sequence data used in this study is deposited in the ENA or SRA, with full accession numbers available in tables S1, S2 and S3. The fastq data for our MinIon assembled genome is available at PRJNA422975.

### Authors contributions

Study conceived by ZZ and AM. Data generated by ZZ and YF. Data analysed by ZZ, SF, CC, YF, and AM. Paper written by ZZ and AM. All authors edited and approved the final manuscript.

